# Mining the heparinome for cryptic antimicrobial peptides that selectively kill gram-negative bacteria

**DOI:** 10.1101/2023.10.19.563059

**Authors:** Daniel Sandín, Javier Valle, Jordi Gómez, Laura Comas, María Nieves Larrosa, Juan José González-López, María Ángeles Jiménez, David Andreu, Marc Torrent

## Abstract

Glycosaminoglycan (GAG)-binding proteins regulating essential processes such as cell growth and migration are essential for cell homeostasis. As both GAGs and the lipid A disaccharide core of gram-negative bacteria contain negatively charged disaccharide units, we hypothesized that GAG-binding proteins could also recognize LPS and enclose cryptic antibiotic motifs. Here, we report novel antimicrobial peptides (AMPs) derived from heparin-binding proteins (HBPs), with specific activity against gram-negative bacteria and high LPS binding. We used computational tools to locate antimicrobial regions in 82% of HBPs, most of those colocalizing with putative heparin binding sites. To validate these results, we synthesized five candidates [HBP1-5] that showed remarkable activity against gram-negative bacteria, as well as a strong correlation between heparin and LPS binding. Structural characterization of these AMPs shows that heparin or LPS recognition promotes a conformational arrangement that favors binding. Among all analogs, HBP-5 displayed the highest affinity for both heparin and LPS, with antimicrobial activities against gram-negative bacteria at the nanomolar range. These results suggest that GAG-binding proteins are involved in LPS recognition, which allows them to act also as antimicrobial proteins. Some of the peptides reported here, particularly HBP-5, constitute a new class of AMPs with specific activity against gram-negative bacteria.

## INTRODUCTION

Glycosaminoglycan (GAG)-binding proteins are a heterogeneous group of proteins mostly associated with the cell surface and the extracellular matrix[1]. They mediate a plethora of functions including signaling, cell proliferation, and coagulation[2–4]. Up to date, most studies of the GAG interactome have focused on protein interactions with heparin, a highly sulfated form of heparan sulfate, due to the commercial availability of heparin and heparin-Sepharose[5]. This has allowed defining the heparin interactome, a highly interconnected network of proteins functionally linked to physiological and pathological processes[6]. Although the structural nature of these proteins is diverse, they share common features, such as the presence of certain domains and motifs[7]. In particular, the CPC’ clip motif is the major contributor to the attachment of heparin (and other sulfated GAGs) to GAG-binding proteins[8]. The motif involves two cationic (Arg or Lys) and one polar (Asn, Gln, Thr, Tyr or Ser, more rarely Arg or Lys) residues with conserved distances between the α carbons and the side-chain center of gravity, defining a clip-like structure where heparin is lodged[9]. The CPC’ clip motif is conserved among all HBPs deposited in the PDB and can be found in many proteins with reported heparin-binding capacity[9].

Recently, we showed that negatively charged polysaccharide-containing polymers, such as heparin and lipopolysaccharides (LPS), can compete for similar binding sites in peptides, and that the CPC’ clip motif is essential to bind both ligands[10]. Our results provide a structural framework to explain why these polymers can cross-interact with the same proteins and peptides and thus contribute to the regulation of apparently unrelated processes in the body. A paradigmatic example is FhuA, an E. coli transmembrane protein involved in the transport of antibiotics such as albomycin and rifamycin[11]. FhuA can bind glucosamine phosphate groups in LPS[12], and we confirmed that a short peptide (YI12WF) retaining most of the LPS-binding affinity of the original protein can also bind heparin with high affinity. When the CPC’ residues in these peptides are mutated, heparin- and LPS-binding activities are largely lost, proving the motif as essential for both ligands. Heinzelmann & Bosshart also showed that human lipopolysaccharide-binding protein (hLBP) can bind heparin and enhance the proinflammatory responses to LPS of blood monocytes[13]. Again, the crystal structure of hLBP bound to N-acetyl-D-glucosamine shows a CPC’ clip motif that could potentially bind heparin. Such observations may prove generalizable to other LPS-binding proteins and may reveal a biological interplay between LPS and heparin. Whether the reverse is true –i.e., HBPs playing a role in LPS binding and potentially in antimicrobial activity– is currently unknown.

Here we show that HBPs contain cryptic AMPs that overlap with heparin-binding regions containing a CPC’ motif. These AMPs show strong selective antimicrobial activity for gramnegative bacteria. They also bind heparin and LPS with high affinity and disrupt the bacterial cell wall. Our results suggest that LPS and heparin bind similar regions in proteins, provided they contain a CPC’ clip motif. HBPs therefore represent a source for new antimicrobials effective against antibiotic-resistant pathogens.

## RESULTS

### Linking heparin affinity and antimicrobial activity

Despite the differences between GAGs and LPS, both contain negatively charged disaccharides in their structure. GAGs are polymers based on variably sulfated repeating disaccharide units. For example, the most common form of heparin is a sulfated disaccharide composed of iduronic acid and glucosamine linked through a β (1→4) bond [IdoA(2S)-GlcNS(6S); Figure 1A]. For its part, LPS is composed of a polysaccharide antigen linked to a lipid A molecule, which is, in turn, a phosphorylated glucosamine (GlcN) disaccharide decorated with multiple fatty acids. The two GlcN units are linked by a β (1→6) bond, and normally contain one phosphate group each (Figure 1B). Based on these structural similarities, we hypothesized that HBPs could also potentially bind the phosphorylated GlcN units of LPS. As heparin-binding sites are commonly associated with short sequential motifs, we reasoned that specific short regions in HBPs could behave as AMPs, binding first to LPS and later destabilizing the outer cell wall and the bacterial membranes.

**Figure 1.**
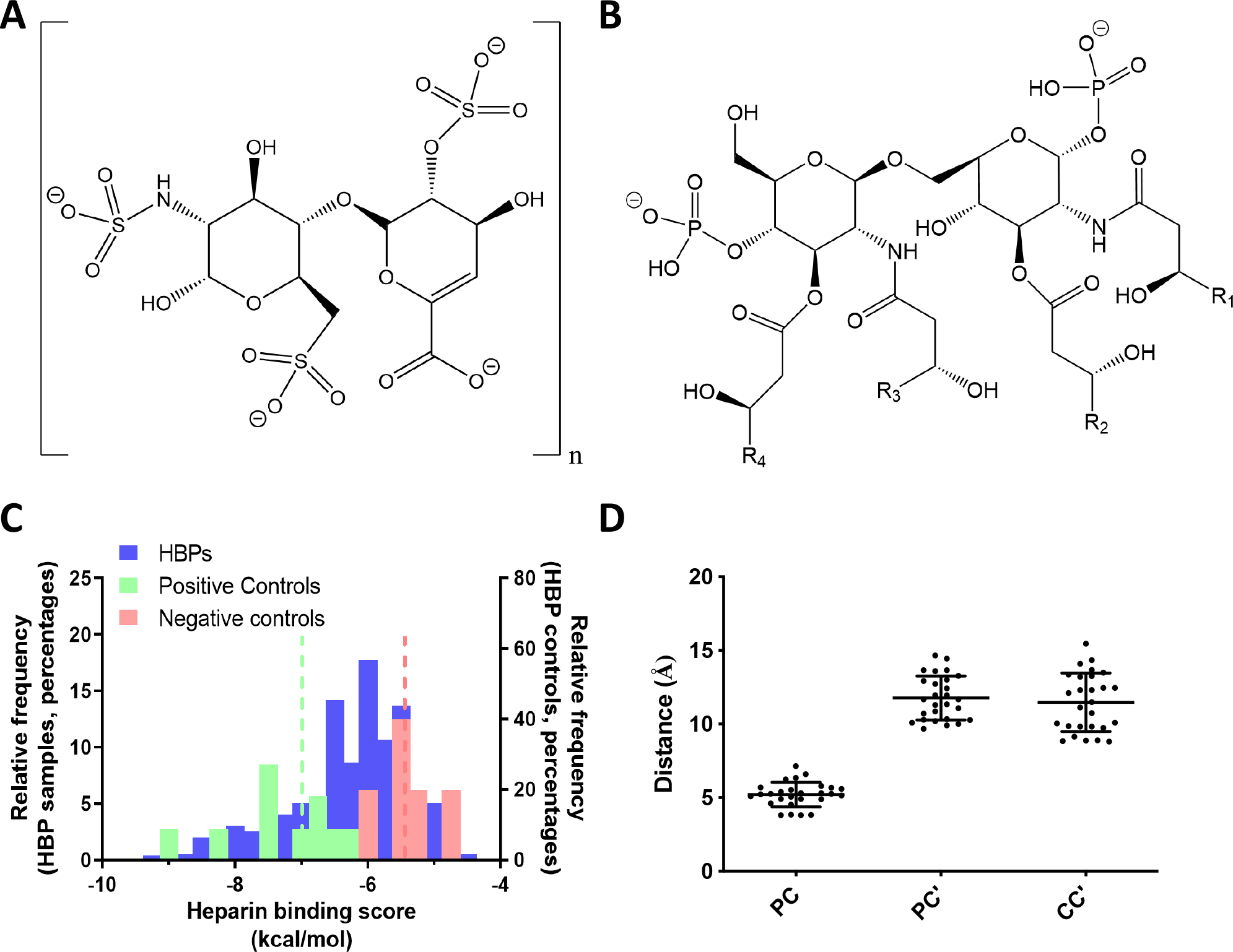
Antimicrobial and heparin binding affinity of HBPs. Structure of **(A)** heparin disaccharide and **(B)** lipid A disaccharide central axis. **(C)** Affinity score distribution of AMPs (blue), positive controls (green, dotted line in the green refers to their mean, -7.0±1.1 kcal/mol) and negative controls (red, dotted line in red refers to their mean, -5.4±0.5 kcal/mol). **(D)** Distances between cationic and polar residues in the best candidates with CPC’ motifs detected. Reference values for PC, PC’ and CC’ residues in CPC’ motifs are 6.0±1.9 Å for PC, 11.6±1.6 Å for PC’ and 11.4±2.4 Å for CC’^7^.

To validate our hypothesis, we inspected all reported HBPs (Supplementary File 1) using the AntiMicrobial Peptide Analyzer (AMPA), a prediction algorithm that can detect the presence of cryptic antimicrobial segments in proteins[14]. Using the default parameters, AMPA detected potential antimicrobial regions in 82% of the HBP set, suggesting that most HBPs contain cryptic AMPs that can be mined by AMPA. According to our hypothesis, these regions should colocalize with heparin-binding sites in HBPs. To ascertain whether the AMPA-retrieved cryptic AMPs could indeed bind GAGs, we first resorted to molecular docking. In AutoDock Vina, a docking region (grid) centered on the antimicrobial segment detected by AMPA was defined and docked with a heparin disaccharide I-S (H1S, α-ΔUA-2S-[1→4]-GlcNS-6S). Results show 76% of the cryptic antimicrobial regions as potential binders of H1S, with affinity comparable to well-defined heparin-binding motifs (Figure 1C). We also examined the presence of CPC’ clips in HBPs with a docking score higher than the average energy calculated for experimentally validated HBPs (-6.8 kcal/mol, 30 proteins) and found that 74% of such regions contain a CPC’ motif with geometric distances compatible with GAG anchoring (Figure 1D). We therefore concluded that heparin-binding regions significantly overlap with cryptic antimicrobial regions in HBPs, hence structural co-localization of antimicrobial activity and GAG recognition can be posited.

### Synthesis and validation of cryptic AMPs from HBPs

To confirm our hypothesis, we synthesized five peptides reproducing the regions with highest AMPA score that also contained a CPC’ clip motif (Table 1, Supplementary Table S1). We used first affinity chromatography to check whether the peptides were able to bind heparin, hence proving that the binding region had been successfully delimited. Indeed, we found the retention times for all peptides in a heparin column to be higher than control, antimicrobial peptide LL-37 (Table 1). In two cases, **HBP-4** and **HBP-5**, affinity was so high that up to 98% buffer B had to be used to dislodge them from the column. So, we could safely conclude that all peptides showed medium-to-strong heparin binding evoking that of parental HBPs.

**Table 1.**
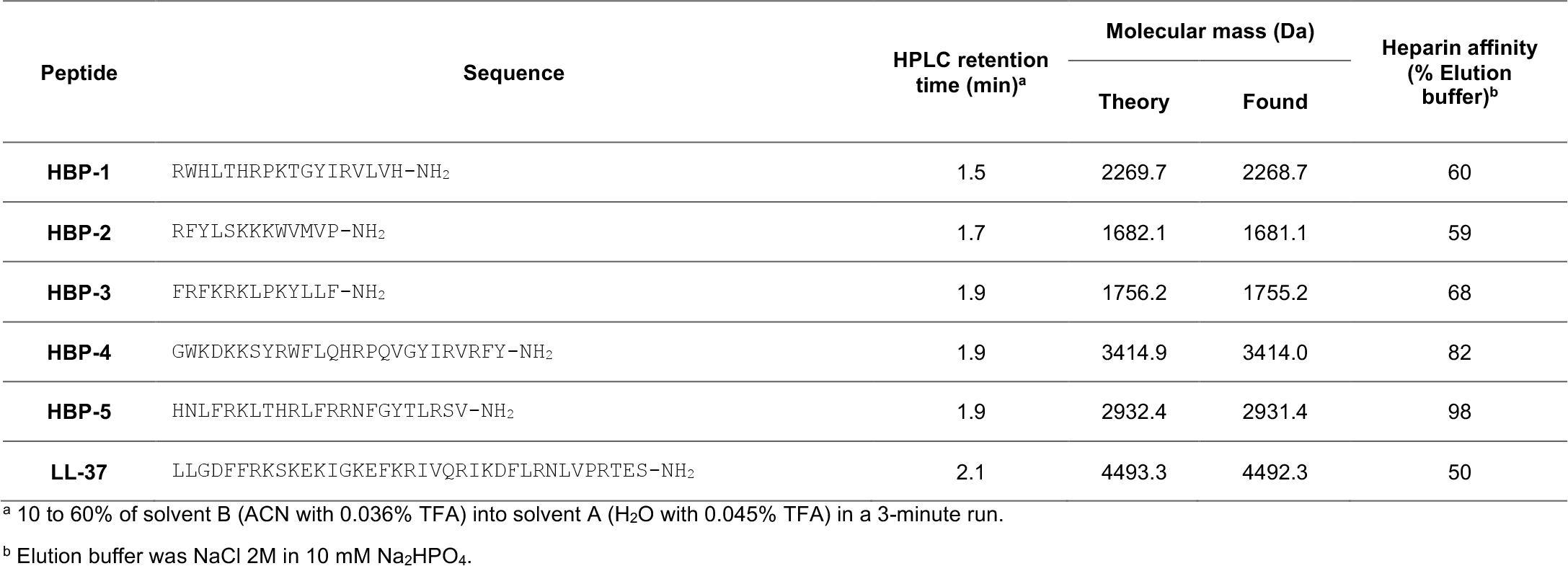
Synthetic peptide analytical data and heparin affinities.

We next inspected antimicrobial activity. Minimal inhibitory concentration (MIC) and minimal bactericidal concentration (MBC) were determined on a panel of gram-negative and gram-positive bacteria. The synthetic peptides displayed strong activity against gram-negative (*Escherichia coli, Acinetobacter baumannii*, and *Pseudomonas aeruginosa*) while being much less active against gram-positive bacteria (*Staphylococcus aureus, Enterococcus faecium*, and *Listeria monocytogenes;* Table 2). This observation is consistent with our hypothesis that, lacking LPS, gram-positives are much less susceptible than -negatives to AMPs. Also, in tune with the hypothesis, peptides with the strongest affinity for heparin (**HBP-4** and **HBP-5)** had the best antimicrobial activity, correlating both observations. In contrast **HBP-2**, the peptide with the lowest affinity for heparin, did not show any significant difference in activity between gram-positive and -negative bacteria, except for *S. aureus*. Antimicrobial activity was also retained against clinical isolates of gram-negative strains (Supplementary Table S2). Specifically, **HBP-4** and **HBP-5** were remarkably active, including multidrug-resistant *P. aeruginosa* strains. Given these encouraging results, we inspected the hemolytic capacity of the peptides as a benchmark of their therapeutic potential as antimicrobials (Supplementary Table S3). Erythrocyte lysis was low for all peptides; only 15% was observed up to 125 μM peptide, in contrast to >30% lysis for LL-37 at the same concentration. On mammalian (MRC-5 and HepGS) cells, similarly favorable results were again found. For **HBP-4**, the (relatively) more cytotoxic peptide, LC_50_ was comparable to LL-37, but **HBP-5** was significantly better. Interestingly, another peptide called NLF20, isolated from the same region of heparin cofactor 2, also displays strong antimicrobial properties[15–17]. Overall, **HBP-5** emerges as the most attractive analog, with a selectivity ratio (LC_50_/MIC) between 50 and 800 (depending on bacterial strain) that must be regarded as outstanding for AMPs and that confirms the hypothesis that HBPs contain cryptic AMPs.

**Table 2.** MIC and MBC data (MIC/MBC, μM) for all peptides against reference strains.

|  | *E. coli* | *A. baumannii* | *Pseudomonas sp* | *S. aureus* | *E. faecium* | *L. monocytogenes* |
| --- | --- | --- | --- | --- | --- | --- |
| **HBP-1** | 1.6 / 1.6 | 1.6 / 1.6 | 3.1 / 3.1 | 50 / >100 | $\geq 100$ / >100 | 6.3 / 12.5 |
| **HBP-2** | 12.5 / 25 | 50 / 50 | 25 / 50 | >100 / >100 | >100 / >100 | 37.5 / >100 |
| **HBP-3** | 3.1 / 3.1 | 0.8 / 0.8 | 6.3 / 6.3 | 25 / 50 | >100 / >100 | 6.3 / 12.5 |
| **HBP-4** | 0.2 / 0.2 | 0.8 / 0.8 | 0.8 / 0.8 | 3.1 / 12.5 | 12.5 / 50 | 1.6 / 3.1 |
| **HBP-5** | 0.4 / 0.4 | 0.2 / 0.2 | 0.8 / 0.8 | 6.3 / 6.3 | 25 / 25 | 1.6 / 3.1 |
| **LL-37** | 1.6 / 3.1 | 6.3 / 6.3 | 0.8 / 0.8 | 25 / 25 | 0.8 / 3.1 | 0.8 / 3.1 |

### Mechanism of action

Given the interesting antimicrobial profiles of HBPs, we investigated their mechanism of action to determine if activity could be related to the interaction with LPS, hence with the cell wall. First, we analyzed LPS binding affinity with the BODIPY-cadaverine assay. The peptides with best antimicrobial activity, **HBP-4** and **HBP-5**, also exhibited strongest LPS binding, comparable to LL-37, while the remaining analogs showed moderate binding, **HBP-2** being the poorest one, again in tune with low antimicrobial activity (Figure 2A, Supplementary Table S4). Consistently, peptide NLF20 was also shown able to disrupt lipopolysaccharide aggregates[18]. This correlation between heparin and LPS affinities strongly suggests that both activities are related (Supplementary Figure S1). The results are also consistent with the lethality curves measured in *E. coli*, in which **HBP-4** and **HBP-5** are fast acting, even more than LL-37, while **HBP-1** and **HBP-2** are the slowest ones (Figure 2B). All peptides showed membrane depolarization abilities comparable to LL-37, according to the DiSC_3_(5) assay (Figure 2C, Supplementary Table S4), with **HBP-5** again scoring highest and **HBP-2** lowest among all analogs. Finally, to directly observe cell wall damage, the morphology of peptide-incubated *E. coli* cells was observed by scanning electron microscopy. In all cases we could detect a clear disruption of the bacterial envelope (Figure 2D), confirming that the peptides act at the outer membrane level, disrupting cell structure and promoting depolarization, eventually resulting in cell death.

**Figure 2.**
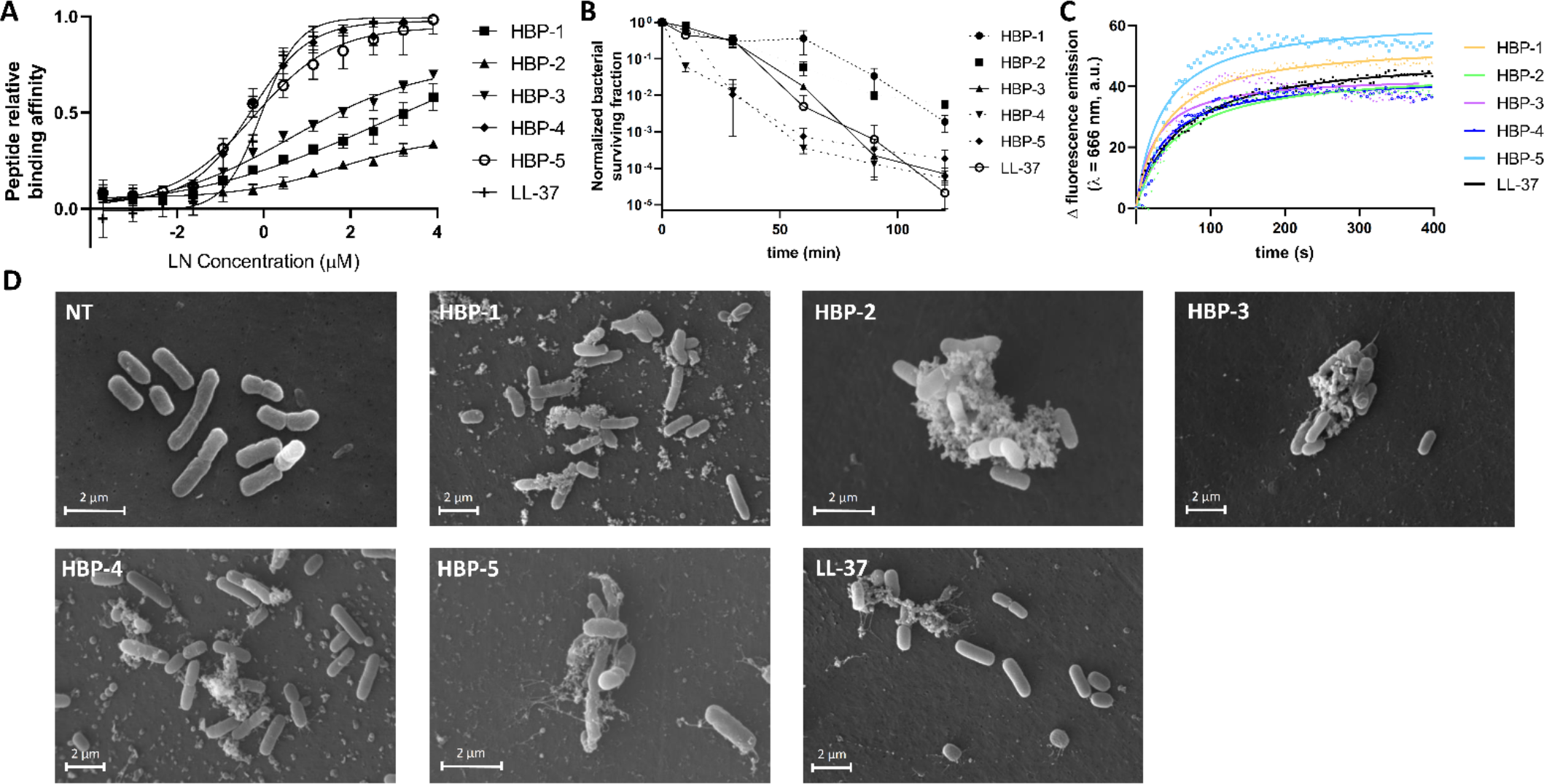
Mechanism of action of HBP-derived antimicrobial peptides. **(A)** LPS affinity measured as an increase in fluorescence emission (λ_em_=620 nm) of BODIPY-cadaverine at different peptide concentrations. **(B)** Bactericidal activity kinetics obtained by treating E. coli planktonic cultures with HBPs. Peptide concentrations were 50 μM HBP-1, 150 μM HBP-2, 50 μM HBP-3, 25 μM HBP-4, 6.3 μM HBP-5 and 50 μM LL-37. (C) Cell depolarization measured as DiSC3(5) fluorescence emission increase after incubating E. coli cells with HBPs. LL-37 was used as positive control of membrane depolarization. All peptides were tested at 10 μM, except HBP-2 at 20 μM. **(D)** SEM pictures of E. coli cells treated with HBPs at same concentrations used in C, after 2h incubation at 37ºC. NT= non peptide control. All measurements were performed in triplicate.

### Structural characterization

To investigate any structural changes occurring upon interaction of the peptides with heparin or cell membranes, we obtained circular dichroism (CD) spectra in buffer, SDS, LPS, and heparin (Figure 3A, Supplementary Tables S5-8). In almost all cases, the structures in water were random, with minima at ∼200 nm. In the presence of SDS, peptides **HBP-3** and **HBP-5** displayed minima near 208 and 222 nm, with a positive band at ∼190 nm, evidencing a shift towards helical conformation. For peptides **HBP1, HBP-2**, and **HBP-4**, a shift towards a minimum at 218 nm was observed, suggesting a beta strand structure. This behavior is typical of AMPs; the random-to-structure transition favors partial insertion into the membrane, promoting depolarization. With LPS, again a transition from random to either helix or beta strand was observed for **HBP-4** and **HBP-5**, less pronounced for the other analogs. This behavior was repeated for all peptides in the presence of heparin, except for **HBP-2**, which remained in disordered conformation. These results are consistent with the above antimicrobial and LPS binding assays in suggesting that LPS and heparin binding triggers a structural arrangement into a more defined, antimicrobially effective structure which, in all cases, is similar to that adopted by the peptide segment in the corresponding original protein (Figure 3A).

**Figure 3.**
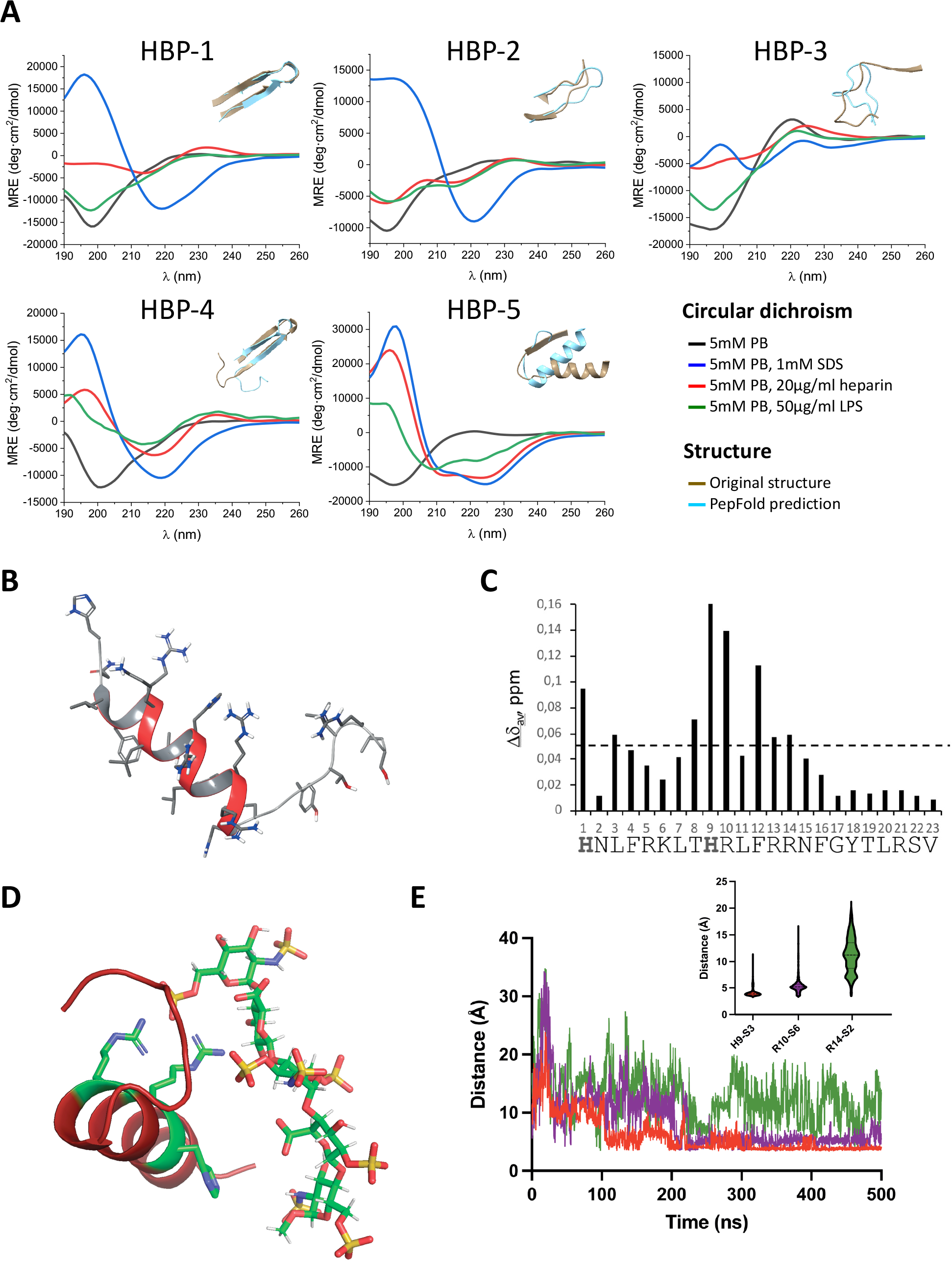
Structural characterization of HBP-5 in different conditions. **(A)** Circular dichroism spectra of HBPs in PBS 5 mM (black lines); PBS 5 mM and SDS 1 mM (blue lines); PBS 5 mM and heparin 20 μg/mL (red lines), and PBS 5 mM and LPS 50 μg/ml (green lines). Overlapping original peptide structures in native protein (in light blue) and PepFold predicted structures (in brown) are added for each peptide in the upper-right corner of each plot. **(B)** Structure of the peptide in DPC micelles as solved by NMR. **(C)** Weighted chemical shift differences (Δδ_w_ = [(δ_HN_^bound^ – δ_HN_^free^)^2^ + (δ_Ha_^bound^ – δ_Ha_^free^)^2^]^1/2^, ppm; see methods) induced by the presence of the heparin disaccharide H1S plotted as a function of peptide sequence. Peptide/disaccharide ratio 1:1. The horizontal line indicates the Δδ_w_ = 0.05 ppm; Residues with values below this line are considered unaffected by interaction; and **(D)** structure of the peptide bound to the heparin analog Arixtra as defined by MD simulation. **(E)** Distances between the CPC’ residues involved in heparin binding as observed in the MD simulation and summary of calculated distances.

As **HBP-5** was the most interesting analog in terms of antimicrobial activity and heparin-binding, we decided to inspect its solution structure by NMR in (i) water, (ii) DPC micelles, and (iii) in the presence of heparin analogs. After assigning ^1^H and ^13^C chemical shifts, we performed a qualitative analysis of the Δδ_Hα_ and Δδ_Cα_ conformational shifts (Δδ = δ^observed^ – δ^random coil^, ppm; see Methods and Supplementary Figure S2). The fact that Δδ_Hα_ and Δδ_Cα_ values are within the random coil range indicates that **HBP-5** is mainly disordered in aqueous solution, as observed previously by CD. In DPC micelles, the stretches of negative Dd_Ha_ and positive Dd_Ca_ values indicate the presence of a highly populated helix structure spanning approximately residues 3-11 (Supplementary Table S9). Structure calculation, which includes medium and long-range distance restraints derived from the observed NOE cross-peaks (see Methods and Supplementary Table S10), showed a well-defined N-terminal helix spanning residues 3-15, three-residues longer than deduced from qualitative analysis of Δδ_Hα_ and Δδ_Cα_ values, and a less ordered non-regular turn-like motif involving residues 18-22 (Figure 3B). The relative arrangement of the a-helix and the turn-like motif is poorly defined. Unfortunately, attempts to retrieve an NMR 3D structure of **HBP-5** complexed with heparin analogs were unsuccessful. Spectra of **HBP-5** with either the fondaparinux (Arixtra^®^) pentasaccharide or the simpler H1S disaccharide acting as heparin analogs did not provide any NOE cross-peaks evidencing intermolecular peptide-sugar contacts, mostly due to the substantial sample precipitation observed, particularly for the fondaparinux pentasaccharide. In the presence of an equimolar amount of H1S disaccharide, many cross-peaks are shifted relative to free **HBP-5** (see Supplementary Figure S3). To identify which residues are most affected upon disaccharide interaction, weighted chemical shift differences (Δδ_w_, ppm) were plotted as a function of peptide sequence (Figure 3C). It is clear in this plot that significant differences are mainly located at the central section, residues 8-14 (Figure 3C). In view of this, and to obtain additional insights into heparin binding of **HBP-5**, we performed a molecular dynamics simulation with fondaparinux. The results show that the pentasaccharide remains in contact with the peptide all along the simulation time, suggesting strong binding. Specifically, we observed a persistent salt bridge between the Arg13 side chain and the S6 sulfate group of the pentasaccharide (Figure 3D). Another salt bridge between Arg10 and the S6 sulfate, plus a loosely defined hydrogen bond between His9 and the S3 were also identified. These three residues (His9, Arg10, and Arg14) form a CPC’ motif with their relative distances maintained throughout the simulation (Figure 3E, Supplementary Figure S4), altogether suggesting a CPC’ clip as a relevant binding element.

## DISCUSSION

Inflammation and coagulation are closely related, with inflammatory proteins often interacting with GAGs and influencing anticoagulant activity[19]. Some proteins play important roles in both processes, such as histidine-rich glycoprotein, an adaptor protein released by platelets, that regulates angiogenesis, immunity, and coagulation[20].

Many proteins, particularly those involved in host defense, can act as reservoirs of AMPs, silently embedded in these protein sequences but produced on demand by host proteases during events such as inflammation, coagulation, etc[21–23]. After a wound, processes to prevent bleeding, remove damaged tissue and keep the lesion free from pathogen entry and subsequent infection are called for[24]. In such scenarios, proteases hydrolyzing surrounding proteins and releasing (formerly) cryptic AMPs to achieve preventive antimicrobial action can play a crucial role. A relevant example is thrombin. While the whole protein does not display antimicrobial activity *per se*, after cleavage its C-terminus displays strong and broad activity[25]. It is therefore not surprising that proteins involved in GAG binding can become an important source of AMPs, hence contribute to preventing infection. This dual action, GAG binding and antimicrobial activity, can be interpreted in structural terms by the similarities between GAG and lipid A structures, both containing negatively charged disaccharide units. It is thus reasonable to suggest that the ability to bind GAGs could also foster LPS recognition, hence allow interaction with the outer membrane of gram-negative bacteria. The fact that LL-37 contains an XBBXBX-motif but lacks strong heparin-binding affinity (Table 1) and fails to show a significant preference for gram-negative bacteria (Table 2), suggests that a CPC’ motif may be relevant to bind both LPS and heparin, as suggested by our structural analysis (Figure 1). Hence, heparin-binding and antimicrobial activity can be related, as previously suggested, due to similar amino acid composition, but a structural arrangement is clearly required to bind to LPS[16,26].

Here, we have shown that GAG-binding proteins can be a source of new AMPs, some with remarkable activity. The fact that these peptides can bind to both heparin and LPS is consistent with the above structural similarity hypothesis, and with the fact that these peptides have much higher activity on gram-negative bacteria. With further optimization, HBP-derived cryptic AMPs should prove useful for treating infections by gram-negative bacteria that are resistant to classic antibiotics and pose huge risks for hospitalized patients.

## Acknowledgements

Financial support from the Spanish Ministry of Economy and Innovation (PDC2021-121544-I00 funded by MCIN/AEI /10.13039/501100011033 and European Union Next GenerationEU/ PRTR to M.T.; project PID2020-114627RB-I00 funded by MCIN/AEI /10.13039/501100011033 to M.T.; PID2020-112821GB-I00 funded by MCIN/AEI /10.13039/501100011033 to M.A.J and AGL2017-84097-C2-2-R to D.A.), and from La Caixa Banking Foundation (HR17-00409, to D.A.). This work has been co-financed by the Spanish Ministry of Science and Innovation with funds from the European Union NextGenerationEU, from the Recovery, Transformation and Resilience Plan (PRTR-C17.I1) and from the Autonomous Community of Catalonia within the framework of the Biotechnology Plan Applied to Health. Work at UPF was funded by the MCI María de Maeztu network of Units of Excellence. D.S. is recipient of pre-doctoral a FPI scholarship (PRE2018-083243) from the Spanish Ministerio de Ciencia e Innovación.

