## Supplementary Information for "Mining the heparinome for cryptic antimicrobial peptides that selectively kill gram-negative bacteria"

### 1. Materials and methods

***Materials.*** *Escherichia coli* BW25113 was obtained from the Coli Genetic Stock Center. *Acinetobacter baumannii* (ATCC 15308), *Pseudomonas sp.* (ATCC 15915), *Staphylococcus aureus* (ATCC 12600), *Enterococcus faecium* (ATCC 19434), *Listeria monocytogenes* (ATCC 19112), and *Streptococcus pyogenes* (ATCC 8668) were obtained from CECT (Valencia, Spain). Clinical strains were obtained from the Vall d'Hebron Hospital Microbiology Service (Barcelona, Spain). MRC-5 and HepG2 cell lines were purchased from ATCC. Horse defibrinated red blood, DiSC3(5) (3,3'-dipropylthiadicarbocyanine Iodide), BODIPY® TR cadaverine, and MTT (3-(4,5-dimethylthiazol-2-yl)-2,5-diphenyl tetrazolium bromide) were from Thermo Fisher (Hampshire, England). *E. coli* lipopolysaccharide and heparin were acquired from Merck (Darmstadt, Germany). Heparin disaccharide I-S trisodium salt (H1S) was from TLC Pharmaceutical Standards (Newmarket, ONT) and fondaparinux (Arixtra®) was a gift from the Hospital del Mar (Barcelona, Spain) pharmacy. The deuterated compounds [D38]-DPC (98 %), and $D_2O$ (99.9%) were purchased from Eurisotop (Saint-Aubin, France). The percentages of deuteration are indicated in parentheses.

***Antimicrobial activity prediction and docking.*** The library of heparin-binding proteins was obtained from Ori et al. All sequences were processed with the AMPA antimicrobial peptide predictor (http://tcoffee.crg.cat/apps/ampa) to define the antimicrobial regions. Best candidates according to AMPA score were used for docking studies using AutoDock Vina using heparin disaccharide H1S (α-ΔUA-2S- [1→4]-GlcNS-6S). Grid boxes were adjusted to the regions as delineated by AMPA. To control for significant binding energy values, we docked H1S to the binding regions of proteins with a solved crystal structure containing a heparin analog. The proteins used as positive controls were, angiogenin (4QFJ), heparin lyase I (3IN9), palmitoleoyl-protein carboxylesterase (4UYW), stromal cell-derived factor 1 (2NWG), peptidoglycan recognition protein 1 (3OGX), C-C motif chemokine 5 (1UL4), heparin cofactor 2 (1JMJ), antithrombin III (1SR5), annexin A2 (2HYV), plasma serine protease inhibitor (3DY0), and heparin lyase (2FUT). Crystal structures of non heparin-binding proteins bound to a non-sulfated disaccharide were used as negative controls, i.e., aconitase (7ACN), R-methyltransferase (R30Q), phytase (3ZHC), bifunctional epoxide hydrolase 2 (1EK2) and calpain-3 (6BGP). The proteins included in the HBPs list with the highest affinity score (higher than the average of positive controls) were checked for the presence of CPC' motifs within their sequence using UCSF Chimera.

***Peptide Synthesis.*** Peptides were synthesized as described previously[1] on H-Rink Amide-ChemMatrix resin in a Prelude instrument (Gyros Protein Technologies, Tucson, AZ) running Fmoc solid-phase peptide synthesis (SPPS) protocols. After sequence assembly, the resin-bound peptides were deprotected in TFA/$H_2O$/triisopropylsilane (95:2.5:2.5 v/v), isolated by cold diethyl ether precipitation and centrifugation at 4800 rpm for 5 min, and lyophilized. Purification was performed on a Luna C18 column (21.2 mm x 250 mm, 10 μm; Phenomenex) in a LC-8 preparative RP-HPLC instrument (Shimadzu, Kyoto, Japan) using a linear gradient of solvent B (0.1% TFA in ACN) into A (0.1% TFA in $H_2O$) for 30 min at 25 mL/min flowrate. Peptides prior and after purification were inspected by analytical RP-HPLC and LC-MS. RP-HPLC was performed on a Luna C18 column (4.6 mm x 50 mm, 3 μm; Phenomenex) in an LC-20AD instrument (Shimadzu) using a linear gradient of solvent B (0.036%TFA in ACN) into A (0.045% TFA in $H_2O$) over 15 min at 1 mL/min flowrate. LC-MS was done in an LC-MS 2010EV instrument (Shimadzu) connected to an Aeris Widepore XB-C18 column (4.6 mm x 150 mm, 3.6 μm; Phenomenex) with a liner gradient of solvent B

(0.08% formic acid (FA) in ACN) into solvent A (0.1% FA in $H_2O$) over 15 min at 1 mL/min flowrate. Peptides with the expected mass and >95% HPLC homogeneity were lyophilized and stored at -20 ºC.

***Heparin-binding affinity assay.*** Heparin binding was evaluated by affinity chromatography on a Heparin HP column (Cytiva, Marlborough, MA) linked to an ÄKTA go FPLC instrument (Cytiva, Marlborough, MA). 10 mL of peptide stocks at 10 µM were loaded in the column, previously equilibrated with binding buffer (10 mM sodium phosphate). Peptides were eluted by a linear gradient of elution buffer (10 mM sodium phosphate, 2 M NaCl) into binding buffer. Heparin affinity for each peptide was defined as the percentage of elution buffer at maximum peak intensity.

***Minimum inhibitory concentration (MIC) and minimum bactericidal concentration (MBC).*** Antimicrobial activities were determined by the classical microtiter broth dilution method recommended by the National Committee of Laboratory Safety and Standards (NCLSS), adapted for AMPs[2]. Briefly, overnight bacterial cell cultures were brought to an exponential growth state ($OD_{600}$=0.4) in MH broth and diluted to a final concentration of 5 x $10^5$ CFU/mL. 1:2 peptide serial dilutions were prepared in 96-well polypropylene plates (Greiner, Frickenhausen, Germany), in MH medium containing 0.2% (w/v) of bovine seroalbumin (BSA) and 0.02% glacial acetic acid. Samples were incubated overnight at 37ºC, and 230 rpm, and the MIC was determined as the last peptide concentration without appreciable visual growth. MBC was determined by transferring the content of the wells to Petri plates with Luria Bertani (LB) agar and incubated overnight at 37ºC. The lowest concentration with no colonies was considered the MBC.

***Killing curve assay.*** Peptide antimicrobial activity was also tested by lethality curve in *E. coli* cultures[3]. First, 50 µL of the peptide stock solution were added to 450 µL of an *E. coli* culture at 5 x $10^5$ cfu/mL in a 1.5 mL polypropylene tube to obtain a final concentration of 1x MIC. Samples were then incubated at 37ºC and 600 rpm in an Accutherm microtube shaking incubator (Labnet, Edison, NJ) for 2 h. Samples of 50 µl were taken at several intervals and plated in LB agar. Plates were incubated overnight at 37 ºC and colonies were counted and compared with the initial inoculum to define the percentage of surviving bacteria.

***Hemolytic activity.*** Peptide toxicity was tested in horse erythrocytes[4]. Horse defibrinated blood was washed three times in phosphate buffer saline (PBS), pH 7.2 to remove excess hemoglobin in the supernatant and then diluted 10x in PBS. Then, 50 µL of erythrocytes were added to a 1.5 mL polypropylene tube and incubated with 50 µL of a 1:2 peptide serial dilution. An erythrocyte disruption (ED) control was prepared by adding 50 µL of 0.1% TritonX-100 in PBS instead of the peptides and an intact erythrocyte (IC) control by adding 50 µL of PBS alone. All samples and controls were incubated for 4 h at 37 ºC. Afterwards, samples were centrifuged at 3000 rpm for 3 min and the supernatants were transferred to a polystyrene 96-well plate and inspected for ED by reading the absorbance at 540 nm in a TECAN Spark instrument (Tecan, Männedorf Switzerland). The hemolysis percentage was calculated as:

$$Hemolysis(\%) = \frac{Sample - IC}{ED - IC}$$

***Lipopolysaccharide binding affinity.*** Displacement of fluorescent cadaverine bound to LPS was used to test peptide affinity to LPS[5]. Briefly, 50 µL of 1:2 peptide serial dilutions in 10 mM HEPES were prepared in polystyrene 96-well plates. Then, a previously incubated mixture of 25 µL of 40 µg/mL LPS and 25 µL of 40 µM cadaverine was added to each well. A control without peptides (NP) was prepared by the addition of 50 µL of HEPES buffer, and one without LPS (NL) by adding 25 μL of HEPES instead of LPS. Plates were read for fluorescence in a TECAN Spark instrument (Tecan, Männedorf Switzerland), with 580 nm excitation and 620 nm emission wavelengths, and with 5 and 10 nm slits respectively. The fraction of peptide bound to LPS was calculated as:

$$Binding = \frac{Sample - NP}{NL - NP}$$

***Bacterial membrane depolarization.*** DiSC3(5) lipophilic dye fixation was tested in *E. coli* fresh cultures to measure depolarization, as previously described[6]. In short, 5 mL bacterial suspensions in exponential phase (~0.4 OD) were washed first with 5mL of buffer A (5 mM HEPES, 20 mM glucose, pH 7.2) and later with 5 mL of buffer B (5 mM HEPES, 20 mM glucose, 100 mM KCl, pH 7.2). Then, bacteria were resuspended in buffer B to an OD of 0.05. 1 mL samples were prepared and then DiSC3(5) was added to a final concentration of 0.4 μM. Fluorescence emission was continuously measured in a Varian Cary Eclipse fluorescence spectrometer (Agilent, Santa Clara, California), with excitation at 625 nm (5 nm slit) and emission at 666 nm(10 nm slit). 10 min after dye addition (estimated time required for DiSC3(5) quenching), peptides were added to the samples to a final 10 μM concentration (except HBP-2, tested at 20 μM). Dye release was monitored for at least 5 min.

***Scanning electron microscopy (SEM).*** 1 mL of *E. coli* bacterial suspensions in exponential growth (~0.4 OD) were treated with 10 μM peptides for 2 h. After treatment, treated cells were filtered through a 0.1 μm Nucleopore filter to attach bacteria and later fixed for 2 h at 4ºC in a buffer containing 2.5% glutaraldehyde in 100 mM Na-cacodylate, pH 7.4. Afterward, cells were coated by immersion in 1% osmium tetroxide in Na-cacodylate buffer for 30 min. Samples were rinsed in the same buffer and dehydrated in ethanol with increasing concentrations (once at 30 and 70% and twice at 90 and 100% (v/v)) for 15 min each. The filters were mounted on aluminum stubs and coated with gold-palladium in a sputter coater (K550; Emitech, East Grinsted, UK). Each sample was later inspected at 15 kV accelerating voltage in an EVO MA 10 scanning electron microscope (Zeiss, Oberkochen, Germany).

***Cytotoxicity in mammalian cells.*** Toxicity in MRC-5 and HepG2 cells was tested by the MTT assay, as previously described[21]. Cell lines were maintained in Eagle's minimum essential medium (MEMα) supplemented with 10% fetal bovine serum (FBS). Cells were cultured in 75 $cm^2$ flasks and then transferred to polystyrene 96-well plates, at 3 x $10^4$ cells per well, and incubated overnight for attachment to the well surface. Then culture media was removed and 1:2 peptide serial dilutions in MEMα were added to each well and later incubated for 4 h. After incubation, peptides were removed and 100 μL of 0.5 mg/mL MTT staining solution in MEMα supplemented with 10% FBS was added to cells and incubated for 1.5 h at 37ºC. Formazan crystals in living cells were detected after disruption with 200 μL dimethyl sulfoxide and then the absorbance was measured at 570 nm in a TECAN Spark instrument.

***Circular dichroism.*** CD spectra of peptides were obtained in four different conditions: 5 mM PBS, 5 mM PBS with 1 mM SDS, 5 mM PBS with 20 μg/mL heparin, and PBS 5 mM PBS with 50 μg/mL LPS. Peptides were dissolved in each condition to a final 10 μM concentration. Samples were transferred to a 0.2 mm quartz cuvette (Hellma, Jena, Germany) and analyzed in a Jasco J-815 CD spectropolarimeter (Jasco, Easton, Maryland) in the 260 to 190 nm range. For each sample, 15 spectra were acquired and averaged. Data were processed with the OriginPro analysis software and subsequently analyzed to predict secondary structure with the CDSSTR method[7] available in the Dichroweb online server[8] (http://dichroweb.cryst.bbk.ac.uk/html/home.shtml).

***NMR spectroscopy.*** NMR samples were prepared by dissolving lyophilized peptide HBP-5 at about 1 mM concentration in aqueous solution ($H_2O/D_2O$ 9:1 v/v), in DPC micelles (30 mM [D38]-DPC in $H_2O/D_2O$ 9:1, v/v) or in aqueous solution with the heparin analog Arixtra or the heparin disaccharide H1S (molar ratios 1:1, 1:0.5). pH was measured using a glass micro-electrode and adjusted to 4.4 by addition of NaOD or DCl. Sodium 2,2-dimethyl-2-silapentane-5-sulfonate (DSS) at a 0.1-0.2 mM concentration was added as internal reference for the $^1H$ chemical shifts. A Bruker AVNEO-600 spectrometer (Bruker Biospin, Karlsruhe, Germany) equipped with a cryoprobe was used to record NMR spectra: 1D $^1H$, 2D $^1H,^1H$-DFQ-COSY (double-filtered-quantum phase-sensitive two-dimensional correlated spectroscopy), $^1H,^1H$-TOCSY (total correlated spectroscopy), $^1H,^1H$-NOESY (nuclear Overhauser enhancement spectroscopy), and $^1H$-$^{13}C$-HSQC (heteronuclear single quantum coherence) at $^{13}C$ natural abundance. TOCSY and NOESY mixing times were 60 ms and 150 ms, respectively. Data were processed using the TOPSPIN software (Bruker Biospin, Karlsruhe, Germany). The NMRFAM-SPARKY software[9] was used to analyze the NMR spectra. $^1H$ chemical shifts were assigned by analysis of the 2D homonuclear spectra using the well-established sequential assignment methodology[10], and $^1H$-$^{13}C$-HSQC spectra were analyzed to assign the $^{13}C$ chemical shifts. The assigned chemical shifts have been deposited at the BioMagResBank (http://www.bmrb.wisc.edu) with accession codes BMRB ID: 51732 (HBP-5 in aqueous solution), 51740 (HBP-5/H1S 1:1) and 51767 (HBP-5 in DPC micelles).

The conformational shifts ($Dd_{Ha}$ and $Dd_{Ca}$) were obtained as the differences between the observed chemical shifts and those in random coil (RC) peptides: $Dd_{Ha} = d_{Ha}^{observed} - d_{Ha}^{RC}$, ppm and $Dd_{Ca} = d_{Ca}^{observed} - d_{Ca}^{RC}$, ppm; $d_{Ha}^{RC}$ and $d_{Ca}^{RC}$ were taken from Wishart et al.[11]. Helix populations (Supplementary Table S10) were estimated from the $^1H_\alpha$ and $^{13}C_a$ chemical shifts as previously described[12]. A weighted value for the chemical shift changes ($Dd_w$, ppm) was defined as:

$$Dd_w = [(d_{HN}^{bound} - d_{HN}^{free})^2 + (d_{Ha}^{bound} - d_{Ha}^{free)2}]^{1/2}$$

Considering all **HBP-5** residues (23 in total), the mean $Dd_w$ is 0.05 ppm. Residues with $Dd_w > 0.05$ ppm can be considered as those mostly affected by interaction.

The structure of **HBP-5** in DPC-micelles was calculated using the iterative procedure for automatic NOE assignment integrated in the CYANA 3.98 program[13]. This algorithm consists of seven cycles of combined automated NOE assignment and structure calculation, in which 100 conformers were computed per cycle. The experimental input data comprises the lists of assigned chemical shifts, and integrated NOE cross-peaks present in the 150 ms NOESY spectra, plus the $\phi$ and $\psi$ dihedral angle restraints. The NOE cross-peaks were integrated using the automatic integration subroutine of the NMRFAM-SPARKY software[9]. The TALOSn webserver[14] was used to obtain the dihedral angle restraints from the $^1H$ and $^{13}C$ chemical shifts. The final structure is the ensemble of the 20 lowest target function conformers calculated in the last cycle. These ensembles were visualized and examined by the MOLMOL program[15].

***Molecular dynamics simulations***

MD simulations with or without Arixtra were conducted using GROMACS v2022.3. The Glycan Reader & Modeler from CHARMM-GUI was used to prepare the system, obtaining the topology and parameter files. The force field CHARMM36 was employed for the protein and Arixtra parameters. Initial structures were solvated in a rectangular box of TIP3P water with a minimum distance of 1.0 nm between protein and the faces of the box. $K^+$ and $Cl^-$ ions were added to neutralize the system at an ionic strength of 0.15 M. Electrostatic interactions were calculated using the particle mesh Ewald method under periodic boundary conditions. Structures were energy-minimized and equilibrated by molecular dynamics for 130 ps. Production simulations were run on a GPU (NVIDIA GeForce RTX 3080 Ti) and 16 CPUs (Intel® Xeon® Gold 6226R CPU @ 2.90GHz) for 500 ns with a time step of 2 fs. NPT conditions were stabilized at 306 K by a V-rescale thermostat[16], and at 1 atm by a Parinello–Rahman barostat[17]. Bonds were constrained using the LINCS algorithm. Representative structures for different analyses were extracted from trajectories with the GROMACS command "gmx cluster", using the gromos algorithm with a RMSD cutoff of 0.18 nm.

### 3. Supplementary Tables

**Supplementary Table S1.** Additional data for peptides HBP-1 to HBP-5.

| Peptide | Sequence | Source Protein[a] | Protein GO Biological processes |
|---|---|---|---|
| **HBP-1** | RWHLTHRPKTGYIRVLVH | Thrombospondin-2 (P35442) | • Cell adhesion<br>• Negative regulation of angiogenesis<br>• Positive regulation of synapse assembly |
| **HBP-2** | RFYLSKKKWVMVP | Alpha-1-antichymotrypsin (ACT, P01011) | • Acute-phase response<br>• Inflammatory response<br>• Maintenance of gastrointestinal epithelium<br>• Negative regulation of endopeptidase activity<br>• Regulation of lipid metabolic process |
| **HBP-3** | FRFKRKLPKYLLF | Amiloride-sensitive amine oxidase (P19801) | • Amine metabolic process<br>• Cellular response to azide<br>• Cellular response to copper ion<br>• Cellular response to heparin<br>• Cellular response to histamine<br>• Putrescine metabolic process<br>• Response to antibiotic |
| **HBP-4** | GWKDKKSYRWFLQHRPQVGYIRVRFY | Cartilage oligomeric matrix protein (P49747) | • Animal organ morphogenesis<br>• Animal organ senescence<br>• Apoptotic process<br>• Artery morphogenesis<br>• BMP signaling pathway<br>• Bone mineralization<br>• Cartilage homeostasis<br>• Chondrocyte development<br>• Chondrocyte proliferation<br>• Collagen fibril organization<br>• Growth plate cartilage development<br>• Limb development<br>• Multicellular organism aging<br>• Multicellular organism growth<br>• Musculoskeletal movement<br>• Negative regulation of apoptotic process<br>• Negative regulation of hemostasis<br>• Platelet aggregation<br>• Positive regulation of chondrocyte proliferation<br>• Protein homooligomerization<br>• Protein processing<br>• Protein secretion<br>• Regulation of bone mineralization<br>• Regulation of gene expression<br>• Response to unfolded protein<br>• Skeletal system development<br>• Skin development<br>• Tendon development<br>• Vascular associated smooth muscle cell development<br>• Vascular associated smooth muscle contraction |
| **HBP-5** | HNLFRKLTHRLFRRNFGYTLRSV | Heparin cofactor 2 (P05546) | • Blood coagulation<br>• Chemotaxis<br>• Negative regulation of endopeptidase activity |

[a] UniProt database codes added in brackets.

**Supplementary Table S2.** MIC and MBC values (μM) of all peptides against gram-negative clinical isolates.

| Peptide | ***E. coli* CFT073** | ***E. coli* 1166795** | ***P. aeruginosa* 827651** | ***P. aeruginosa* 827632** | ***A. baumannii* 3878** | ***A. baumannii* 3880** |
|---|---|---|---|---|---|---|
| | MIC/MBC | MIC/MBC | MIC/MBC | MIC/MBC | MIC/MBC | MIC/MBC |
| **HBP-1** | 3.1 / 3.1 | 1.6 / 1.6 | 25 / 50 | 25 / 25 | 1.6 / 1.6 | 12.5 / 12.5 |
| **HBP-2** | 50 / 100 | 12.5 / 12.5 | >100 / >100 | >100 / >100 | 50 / 50 | >50 / >50 |
| **HBP-3** | 12.5 / 25 | 1.6 / 1.6 | 12.5 / 12.5 | 25 / 25 | 6.3 / 6.3 | 6.3 / 6.3 |
| **HBP-4** | 0.8 / 1.6 | <0.1 / <0.1 | 3.1 / 6.3 | 6.3 / 6.3 | 0.8 / 0.8 | 1.6 / 1.6 |
| **HBP-5** | 0.4 / 0.8 | <0.1 / <0.1 | 0.8 / 1.6 | 1.6 / 1.6 | 0.2 / 0.2 | 0.4 / 0.4 |

**Supplementary Table S3.** Hemolytic and cytotoxic activities of peptides.

| Peptide | Hemolysis (%)[a] | $LC_{50}$ (MRC-5 cells, μM) | $LC_{50}$ (HepGS cells, μM) |
|---|---|---|---|
| **HBP-1** | 4.6 ± 0.6 | >200 | >200 |
| **HBP-2** | 2 ± 1 | >200 | >200 |
| **HBP-3** | 4 ± 1 | >200 | >200 |
| **HBP-4** | 15.5 ± 0.1 | 35 ± 1 | 38 ± 13 |
| **HBP-5** | 10.3 ± 0.2 | 69 ± 2 | 80 ± 7 |
| **LL-37** | 32.7 ± 0.7 | 26 ± 3 | 53 ± 2 |

[a] Hemolysis data was assayed at a peptide concentration of 125 μM.

**Supplementary Table S4.** $EC_{50}$ and $t_{1/2}$ values for HBP peptides and LL-37 (Figures 3B and 3C respectively).

| Peptide | LPS Affinity $EC_{50}$ (μM) | $DiSC_3(5)$ $t_{1/2}$ (s) |
|---|---|---|
| **HBP-1** | 42 ± 16 | 38 ± 8 |
| **HBP-2** | 1500 ± 600 | 53 ± 12 |
| **HBP-3** | 7 ± 5 | 25 ± 5 |
| **HBP-4** | 0.7 ± 0.6 | 37 ± 3 |
| **HBP-5** | 0.9 ± 0.7 | 34 ± 2 |
| **LL-37** | 0.9 ± 0.8 | 58 ± 3 |

**Supplementary Table S5.** Calculated secondary structure percentages by circular dichroism in 5 mM PB using CDSSTR in dichroweb (http://dichroweb.cryst.bbk.ac.uk/html/process.shtml).

| Peptide | α-Helix | β-Strand | Turns | Unordered | Total |
|---|---|---|---|---|---|
| **HBP-1** | 0.07 | 0.33 | 0.24 | 0.35 | 0.99 |
| **HBP-2** | 0.04 | 0.32 | 0.24 | 0.37 | 0.97 |
| **HBP-3** | 0.09 | 0.31 | 0.26 | 0.34 | 1 |
| **HBP-4** | 0.06 | 0.36 | 0.23 | 0.34 | 0.99 |
| **HBP-5** | 0.12 | 0.27 | 0.26 | 0.34 | 0.99 |

**Supplementary Table S6.** Calculated secondary structure percentages by circular dichroism in 5 mM PB, 1 mM SDS using CDSSTR in dichroweb (http://dichroweb.cryst.bbk.ac.uk/html/process.shtml).

| Peptide | α-Helix | β-Strand | Turns | Unordered | Total |
|---|---|---|---|---|---|
| **HBP-1** | 0.07 | 0.39 | 0.23 | 0.32 | 1.01 |
| **HBP-2** | 0.10 | 0.38 | 0.23 | 0.30 | 1.01 |
| **HBP-3** | 0.11 | 0.35 | 0.25 | 0.30 | 1.01 |
| **HBP-4** | 0.05 | 0.38 | 0.23 | 0.33 | 0.99 |
| **HBP-5** | 0.50 | 0.24 | 0.15 | 0.19 | 1 |

**Supplementary Table S7.** Calculated secondary structure percentages by circular dichroism in 5 mM PB, LMW heparin 20 µg/mL using CDSSTR in dichroweb (http://dichroweb.cryst.bbk.ac.uk/html/process.shtml).

| Peptide | α-Helix | β-Strand | Turns | Unordered | Total |
|---|---|---|---|---|---|
| **HBP-1** | 0.04 | 0.45 | 0.22 | 0.29 | 1 |
| **HBP-2** | 0.02 | 0.43 | 0.22 | 0.32 | 0.99 |
| **HBP-3** | 0.06 | 0.46 | 0.19 | 0.28 | 0.99 |
| **HBP-4** | 0.06 | 0.42 | 0.21 | 0.31 | 1 |
| **HBP-5** | 0.37 | 0.23 | 0.17 | 0.23 | 1 |

**Supplementary Table S8.** Calculated secondary structure percentages by circular dichroism in 5 mM PB, LPS 50 µg/mL using CDSSTR in dichroweb (http://dichroweb.cryst.bbk.ac.uk/html/process.shtml).

| Peptide | α-Helix | β-Strand | Turns | Unordered | Total |
|---|---|---|---|---|---|
| **HBP-1** | 0.02 | 0.39 | 0.23 | 0.35 | 0.99 |
| **HBP-2** | 0.01 | 0.39 | 0.21 | 0.36 | 0.97 |
| **HBP-3** | 0.06 | 0.38 | 0.21 | 0.33 | 0.98 |
| **HBP-4** | 0.06 | 0.44 | 0.2 | 0.3 | 1 |
| **HBP-5** | 0.2 | 0.27 | 0.25 | 0.29 | 1.01 |

**Supplementary Table S9.** Averaged $\Delta\delta_{H\alpha}$ and $\Delta\delta_{C\alpha}$ values for **HBP-5** in aqueous solution at pH 5.0 and in DPC micelles (30 mM DPC) at pH 5.0 at 25ºC. Percentage of helical structure was estimated from these values. [a] Errors are reported as the standard deviation.

| HBP-5 | | | | | | |
|---|---|---|---|---|---|---|
| **Conditions** | **Helical residues** | **$\Delta\delta_{H\alpha}$, ppm** | **% helix from $\Delta\delta_{H\alpha}$** | **$\Delta\delta_{C\alpha}$, ppm** | **% helix from $\Delta\delta_{C\alpha}$,** | **Averaged % helix[a]** |
| Aqueous solution | 3-11 | -0.039 | 10 | +0.02 | 5 | 5±5 |
| DPC micelles | 3-11 | -0.307 | 79 | +2.95 | 96 | 85±9 |

**Supplementary Table S10.** Summary of structural statistic parameters for the ensemble of the 20 lowest target function conformers calculated for **HBP-5** in DPC micelles.

| **HBP-5 in DPC micelles** | |
|---|---|
| **Number of distance restraints** | |
| Intraresidue & sequential (i – j ≤ 1) | 198 |
| Medium range (1 < \|i – j\| < 5) | 37 |
| Long range (\|i – j\| ≥ 5) | 3 |
| Total number | 238 |
| Averaged total number per residue | 10.3 |
| **Number of dihedral angle constraints** | |
| Number of restricted ϕ angles | 21 |
| Number of restricted ψ angles | 20 |
| Total number | 41 |
| **Pairwise RMSD (Å)** | |
| **All residues** | **2-22** |
| Backbone atoms | 1.6±0.7 |
| All heavy atoms | 2.6±0.7 |
| **N-terminal helix** | **3-15** |
| Backbone atoms | 0.3±0.2 |
| All heavy atoms | 1.6±0.3 |
| **Ramachandran plot (%)** | |
| Most favoured regions | 93.8 |
| Additionally allowed regions | 6.2 |
| Generously allowed regions | 0.0 |
| Disallowed regions | 0.0 |

### 4. Supplementary Figures

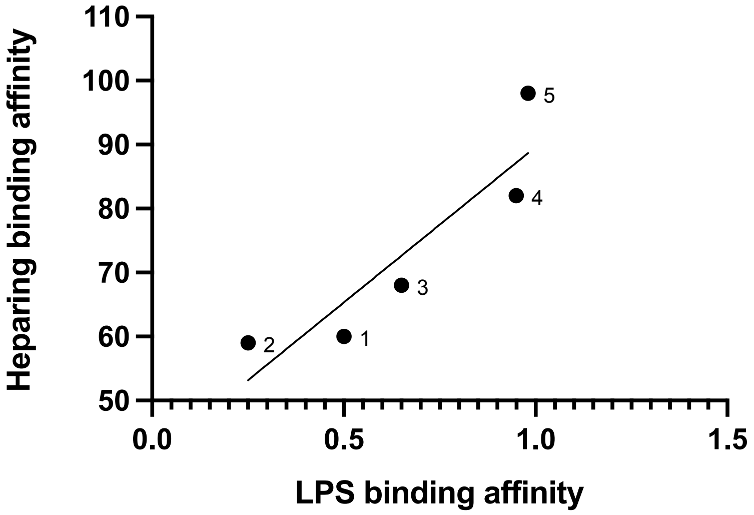


**Supplementary Figure S1. Correlation of heparin-binding activity and LPS-binding activity**. Heparin affinity is measured as the percentage of elution of the peptides in a heparin affinity column and LPS affinity is measured as $EC_{50}$ values as calculated from the BODIPY-cadaverine assay.

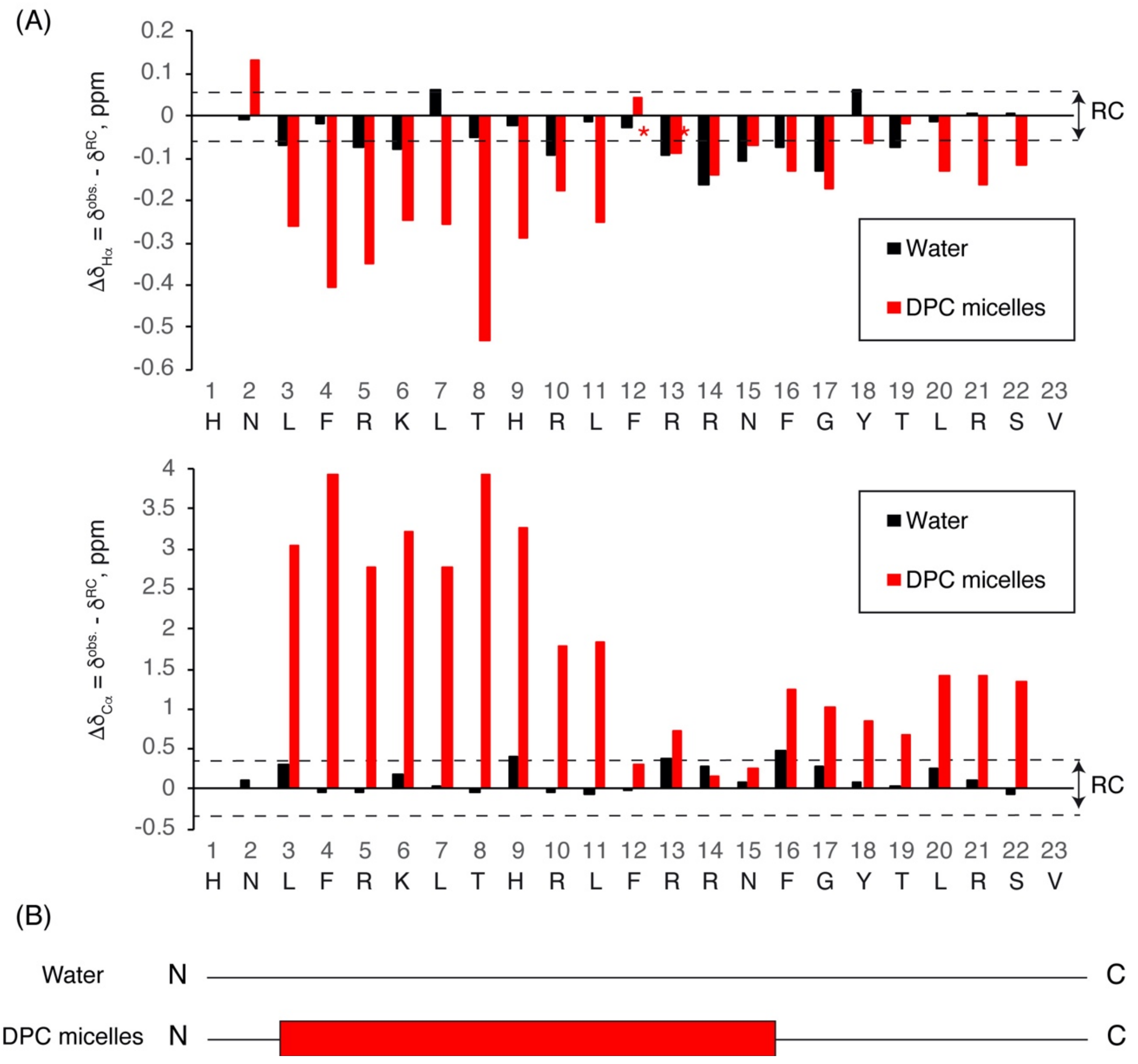


**Supplementary Figure S2.** (A) $Dd_{Ha}$ and $Dd_{Ca}$ conformational shifts for HBP_5 in aqueous solution (black bars) and in DPC micelles (red bars) at pH 5.5 and 25ºC plotted as a function of peptide sequence. The two dashed lines indicate the random coil range (RC). (B) Schematic representation of the structural features in aqueous solution and in DPC micelles. Helices are shown as red rectangles.

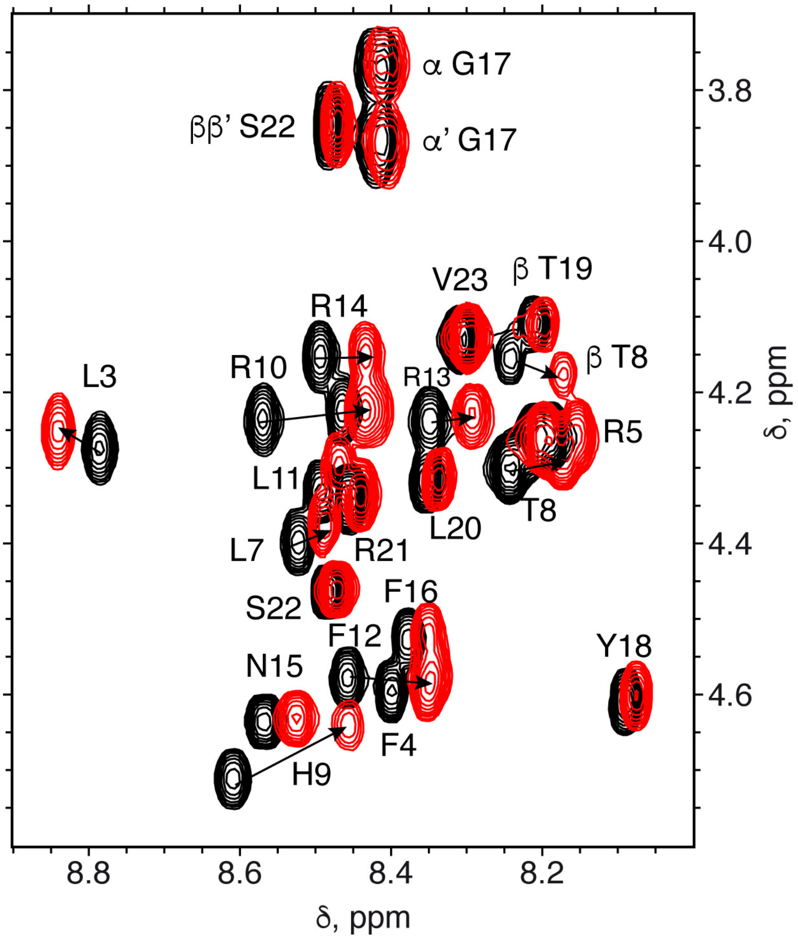


**Supplementary Figure S3.** Overlay of selected regions of 2D $^{1}H$,$^{1}H$ TOCSY spectra for free **HBP-5** (black contours) and for **HBP-5** in the presence of heparin disaccharide HIS at 1:1 ratio (red contours). In both cases, aqueous solution at pH 5.5 and 5ºC. Cross-peaks between a and $H_N$ protons are labelled, and the arrows connect peaks corresponding to free and H1S-bound **HBP-5**.

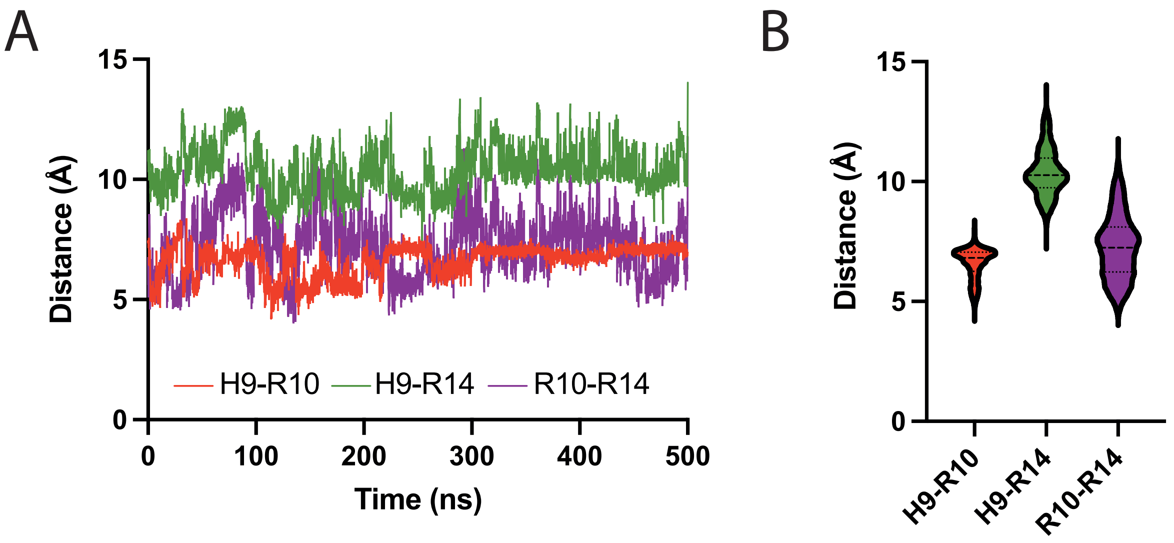


**Supplementary Figure S4. Molecular dynamics simulation of HBP-5 in presence of Arixtra heparin analog**. (A) Distance between residues H9 and R10 (red), H9 and R14 (green), and R10 and R14 (purple) during the simulation. (B) Average distances for each residue pair over the simulation.

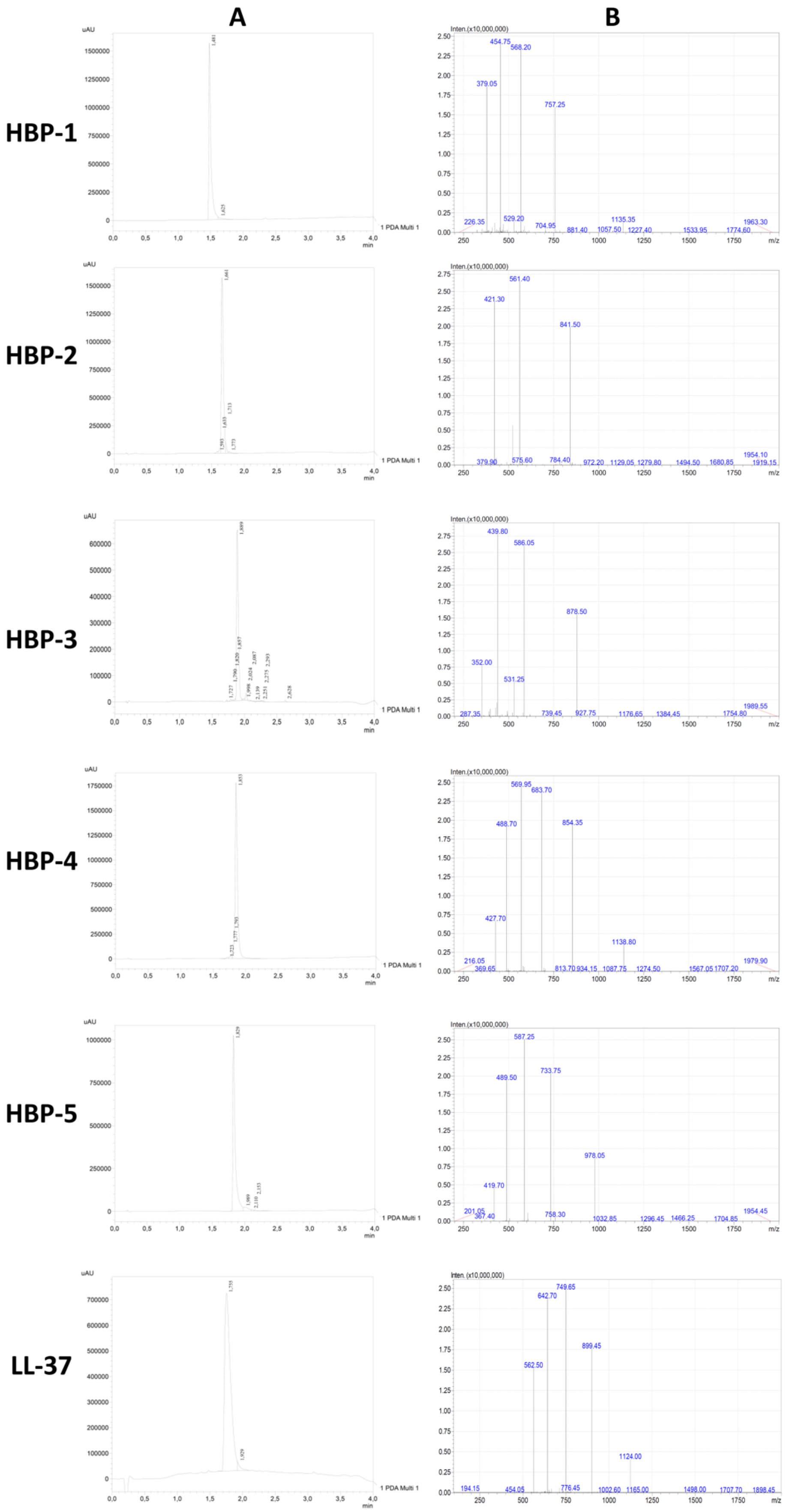


**Supplementary Figure S5.** HPLC chromatograms **(A)** and MS spectra **(B)** of purified synthetic peptides.
